# Melatonin suppression by light involves different retinal photoreceptors in young and older adults

**DOI:** 10.1101/2023.06.02.543372

**Authors:** Raymond P. Najjar, Abhishek S. Prayag, Claude Gronfier

## Abstract

**Introduction:** Age-related sleep and circadian rhythm disturbances may be due to altered non-visual photoreception. Here, we investigated the temporal dynamics of light-induced melatonin suppression in young and older individuals.

**Methods:** In a within-subject design study, young and older participants were exposed for 60 minutes (0030-0130 at night) to 9 narrow-band lights (range: 420 to 620 nm). Plasma melatonin suppression was calculated at 15, 30, 45, and 60 min time intervals. Individual spectral sensitivity of melatonin suppression and photoreceptor contribution were predicted for each interval and age group.

**Results:** In young participants, melanopsin solely drove melatonin suppression at all time intervals, with an invariant peak sensitivity at ∼485 nm established only after 15 minutes of light exposure. Conversely, in older participants, spectral light-driven melatonin suppression was best explained by a model combining melanopsin + L-cones with a stable peak sensitivity (∼499 nm) at 30, 45, and 60 minutes of light exposure.

**Conclusion:** Aging is associated with a distinct photoreceptor contribution to melatonin suppression by light. While in young adults melanopsin-only photoreception is a reliable predictor of melatonin suppression, in older individuals this process is jointly driven by melanopsin and L-cones. These findings offer new prospects for customizing light therapy for older individuals.

## Introduction

In the retina, a small subset of intrinsically-photosensitive retinal ganglion cells (ipRGCs), with a peak sensitivity to blue light (λ_max_ ∼480 nm) (Berson et al., 2002; Dacey et al., 2005), integrate phototransduced light signals from rods (λ_max_ ∼505 nm) and cones (S-cones, λ_max_ ∼420 nm; M-cones, λ_max_ ∼530 nm; L-cones, λ_max_ ∼558 nm) (Dartnall et al., 1983; Spitschan et al., 2019b), before projecting to the SCN and other non-visual brain areas (Gooley et al., 2001; Hattar et al., 2006; Hannibal et al., 2014). This set-up oversees the spectro-temporal sensitivity of various non-visual responses to light including the pupillary light response (PLR), melatonin suppression, and circadian entrainment.

In mammals, including humans, ipRGCs integrate the input from cones and rods in an additive or opponent manner (Dacey et al., 2005; Spitschan et al., 2014, 2019a; Cao et al., 2015; Woelders et al., 2018). In non-human primate retinas, single unit recordings of ipRGCs highlighted that melanopsin-based signalling is combined with a convergent ON input from L- and M-cones, and an OFF input from S-cones in response to 550 nm light (Dacey et al., 2005). Similarly, in humans, intricate combinations of photoreceptoral signalling (melanopsin, S-, M-, L-cones, hereafter described as S, M, and L) have been shown to regulate the PLR and other non-visual responses to light (Spitschan et al., 2014, 2019a; Cao et al., 2015; Woelders et al., 2018). For instance, Spitschan and colleagues suggested that the PLR is governed by an ON contribution from melanopsin and L+M; but an OFF contribution from S (Spitschan et al., 2014). Compared to the PLR, recent findings from our group and others suggest that melanopsin may fully account for melatonin suppression by light (Prayag et al., 2019; Spitschan et al., 2019a; Brown, 2020). Accordingly, a recent analytical paper by Gimenez and colleagues highlighted a sigmoidal dose-response relationship between melatonin suppression and melanopic equivalent daylight illuminance (EDI), but not with photopic illuminance (Giménez et al., 2022). While all of the aforementioned studies focused on healthy young individuals, to date, the impact of aging on photoreceptoral contribution to non-visual responses to light in general, and melatonin suppression in particular, has not been investigated.

Aging is associated with a host of changes, in sleep and circadian physiology, in neuronal and behavioral plasticity, all of which contribute to the neurobehavioral alterations observed across the lifespan (Dijk et al., 2000a; Duffy et al., 2002, 2015; Ohayon et al., 2004; Cajochen et al., 2006b; Li et al., 2006; Skeldon et al., 2016; Mander et al., 2017). These changes could be due, in part, to age-related alterations of the circadian timing system, (Dijk et al., 2000b; Niggemyer et al., 2004; Cajochen et al., 2006a) either at the eye level (Teikari et al., 2012; Najjar et al., 2014, 2016), at the central clock level in the SCN (Cayetanot et al., 2005; Aujard et al., 2006; Hofman and Swaab, 2006; Gibson et al., 2009) or at output levels (Kunz et al., 1998; Kunz, 1999; Claustrat et al., 2005; Kessler et al., 2011). While others have reported reduced melatonin suppression by white (Duffy et al., 2007) and short wavelength light (Herljevic et al., 2005) in older participants, our team showed no reduction in suppression in response to 60 minutes of light exposure, but revealed a shift in the peak spectral sensitivity of melatonin suppression to longer wavelength lights in the elderly compared to young (Najjar et al., 2014). Our findings suggested a re-shuffling of the photoreceptor contribution to melatonin suppression with aging.

In this study, we investigated the temporal dynamics of melatonin suppression in response to light at night, in young and older participants, at the finer timescale of 15-minute intervals. In addition, we further addressed photoreceptoral contribution to melatonin suppression by light over time in both age groups.

## Material and Methods

### Participants

Thirteen healthy participants were recruited in a within subject design study (130 light exposure sessions overall, see details below) *via* local advertisement. Participants were assigned into two age groups, a younger group including five participants (5 men, 25.8 ± 0.73 years, mean ± SE), and an older group including 8 participants (2 men and 6 women, 59.4 ± 0.99 years, mean ± SE). The screening procedure included a comprehensive ophthalmologic examination by a collaborating ophthalmologist that included slit lamp examination, standard automated perimetry (Hymphrey™, Sita-standard 24-2), and colour vision tests (Farnsworth-Munsell 100 Hue test) to exclude patients with ocular diseases. All participants were phakic in both eyes, non-smokers, non-alcoholic and free of medications known to affect sleep, the visual and circadian systems and melatonin secretion. All participants were in good mental health (Mini Mental Score (MMS) > 28) and had no sleep or psychiatric disorders as revealed by the Pittsburgh Sleep Quality Index (PSQI), the SIGH SAD and the PIDS-SA questionnaires. None of the participants was found to be an extreme morning or evening type by the Horne-Östberg morningness-eveningness questionnaire (MEQ). Participants were asked to maintain a regular sleep wake cycle during and three weeks prior to study, and their rest activity pattern was verified by wrist actimetry (Actitrac ™, IM Systems). Participants’ sleep stability was also verified throughout the study using sleep diaries.

Prior to participation, participants provided written informed consent after receiving a detailed description of the procedures and purpose of the experiments. All procedures were in compliance with the institutional guidelines and the Declaration of Helsinki, and the protocol was approved by the Institutional Review Board.

### Experimental protocol

Each participant underwent 10 experimental night sessions (9 light sessions and 1 control session) scheduled between 1930 and 0400 (see Najjar *et al*. 2014 for more details). During each session, participants were exposed for 60 minutes to one of 9 monochromatic light stimuli (peak wavelengths: 420 to 620nm). Conversely, participants were not exposed to light during the dark control session, in order to assess their individual endogenous melatonin profile. The order of the sessions was randomized for each participant, and sessions were separated by a minimum of one week, to disperse any potential phase shifting effect of light on the circadian system from one session to another. Participants were maintained in an ambient very dim light (< 1 lux) between 2000 and 2200 in order to measure their unmasked dim light melatonin onset (DLMO) and in complete darkness (obscure room and blindfolded) from 2200 to 0030 and 0130 to 0400. During that period of time, participants were in semi-recumbent posture (45°) in their bed in order to avoid possible effects of postural changes on melatonin concentration (Deacon and Arendt, 1994). Light exposure was scheduled between 0030 and 0130 for all subjects. During the light stimulation, participants were in a sitting position (90°). Pupils were dilated using tropicamide (2 mg/0.4 ml, Novartis Ophthalmics, Rueil-Malmaison, France), with one drop repeated three times prior to light exposure (−60, -45 and -30 min). Participants were not allowed to use their phones or perform any visual tasks requiring light (reading, watching television, etc.) throughout the session but were allowed to sleep before and after the light stimulation. Fluid intake was controlled to avoid the dilution of melatonin concentrations in blood samples.

Blood samples (4 ml) were collected in a heparinized plastic tube *via* an indwelling catheter at different time intervals during the session (every 60 min from 2000 to 2200 and 0300 to 0400, 30 min from 2000 to 2330 and 0200 to 0300, and 15 min between 2330 and 0200). Following the blood sample taken at 0030, 4 blood samples were taken during the light exposure at 15, 30, 45 and 60 minutes after light onset. Samples were then stored at 4°C and centrifuged at the end of the session (2000 g for 20 minutes). Plasma was decanted and stored at –20°C until assayed. All experimental sessions were performed at the Chronobiology Platform (Edouard Herriot Hospital, Lyon, France).

### Light exposures

A white tungsten halogen light (3250K) was collimated through monochromatic filters (full width at half maximum (FWHM) = ∼10 nm, Omega Optics, λ_max_ 420, 440, 460, 480, 500, 530, 560, 590, and 620 nm). After diffusion by an opaline filter, light was projected into a Ganzfeld sphere with a diameter of 45 cm, coated with white high reflectance paint. The sphere allowed a constant and uniform illumination of the entire retina. The participant’s head was maintained in a constant position inside the sphere by an ophthalmologic head holder. Light exposures were calibrated to be of equal photon density (3.16 x 10^13^ photons/cm^2^/sec) at the subjects’ eye level using a radiometer (International Light ® IL 1700, Peabody, MA, USA). The spectral emission characteristics of the lights were verified using a spectrophotometer (Ocean Optics USB4000, Ocean Optics, Dunedin, FL, USA). Gaze behaviour and pupil dilation were monitored by the experimenter to ensure constancy of exposure for all conditions.

### Melatonin radioimmunoassay

For technical reasons, two radioimmunoassay (RIA) were used in the study. Out of 13 subjects, the samples of 7 participants (5 young and 2 older) were assayed using a radioimmunological method developed by Claustrat and colleagues (Claustrat et al., 1984). Melatonin concentrations were determined in duplicate after diethylether extraction. The assay used an iodinated ligand. Functional sensitivity was 2.6 pg/ml. Inter-assay coefficient of variation was 11% at 50 pg/ml and 13% at 100 pg/ml (n=15). Intra-assay coefficient of variation was 7% at 50 pg/ml and 9% at 100 pg/ml (n=12).

The samples of the remaining 6 subjects were assayed using the Bühlmann melatonin radioimmunoassay kit (Bühlmann AG, Switzerland). With this assay, melatonin concentrations were determined in duplicate after C18 solid phase extraction by a double antibody radioimmunoassay based on the Kennaway G280 anti-melatonin antibody (Vaughan, 1993). Analytical sensitivity was 0.3 pg/ml and functional sensitivity 0.9 pg/ml. Inter-assay coefficient of variation was 18.1% at 1.9 pg/ml and 10.1% at 24 pg/ml (n=18 samples). Intra-assay coefficient of variation was 9.4% at 1.9 pg/ml and 6% at 24 pg/ml (n=18 samples).

The resulting concentrations from the two assays were highly correlated (n = 15 samples from 1 participant; r = 0.90, p<0.001).

### Data and statistical analysis

#### Melatonin secretion and suppression

Individual melatonin amplitude and dim light melatonin onset (DLMO) were calculated as the maximal secretion of the smoothed melatonin profiles (locally weighted scatterplot smoothing LOESS method, span = 0.5) and the 25% upward crossing (100% being the melatonin amplitude calculated between the 0030 and the 2000 concentrations), respectively. DLMOs were compared across sessions and between subjects using a two-way analysis of variance (ANOVA) for repeated measures (within: time; between: subject). DLMO were compared between young and aged groups using a two-way ANOVA for repeated measures with (within: wavelength; between: groups). Melatonin secretion amplitudes assessed during the control session were compared between groups using a Mann-Whitney Rank Sum Test.

The percentage of melatonin suppression was calculated for each participant’s light and control sessions (Eq.1).

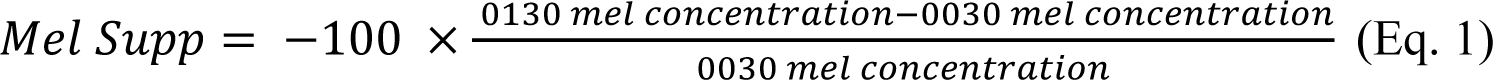

To account for individual differences in melatonin secretion profiles that are independent of light (Gaddy et al., 1993; Brainard et al., 1997), the percent melatonin suppression score was then normalized to the control session (%CA) (Eq. 2).

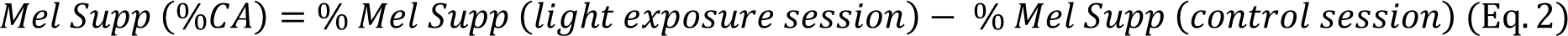

#### Modelling the photoreceptoral contribution to melatonin suppression

We used two approaches to model retinal photoreceptors’ contribution to melatonin suppression in both age groups.

**Approach 1:** Given that melanopsin strongly predicts melatonin suppression by light in the young (Prayag et al., 2019; Brown, 2020), we first modelled a melanopsin-only contribution to melatonin suppression in the young and older groups, at each of the 4 timepoints of light exposure (15, 30, 45 and 60 minutes after light onset). The Govardovskii alpha-band template (Govardovskii et al., 2000) was used to generate opsin templates with constrained peak sensitivity. A melanopsin sensitivity function was fitted on the average, control-adjusted, percentage melatonin suppression, measured at each time interval and for each group (Eq. 3).

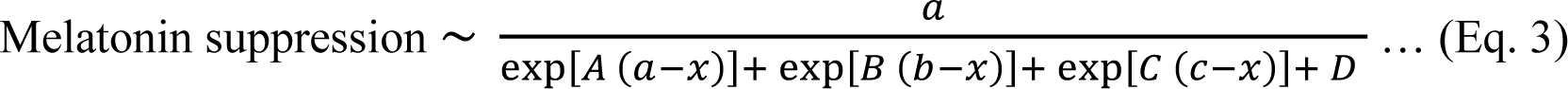

where 𝑥 = 480/*wavelength*. Other parameter values were *A* = 69.7, *a* = 0.88, *B* = 28, *b* = 0.922, *C* = -14.9, *c* = 1.104, *D* = 0.674 as in the Govardovskii et al., (2000).

The amplitude, *a*, was not constrained to the value of 1 as in Govardovskii et al., (2000), but allowed to be optimally determined by the nonlinear least-squares curve-fitting procedure. The peak sensitivity of melanopsin was fixed at 480 nm (CIE, 2018).

Given that the dataset used is identical to the previously published data in Najjar et al., (2014), the lens filtering spectra obtained for the young and the older in that study was applied so as to account for the lens transmittance and its change with aging. Each fitted function in the young and older participants was accounted for pre-receptoral filtering by multiplying the function with its respective ocular lens transmittance spectra, published in Najjar et al., (2014). Peak spectral sensitivities for both groups (λ_max_), at the four time-intervals following light onset, were predicted using this melanopsin nomogram corrected for the respective transmittance of the ocular media in younger and older participants (Table 1). The correlation coefficients (R² and adjusted-R²) were retrieved as a measure of the goodness of each model and for comparison with coefficients previously reported in other studies. The adjusted-R^2^ accounted for the number of predictors in each model. The fitting of the curves was carried out using nonlinear least squares fitting procedures in R v. 3.6.2 statistical environment (http://cran.r-project.org).

**Table 1:** Peak spectral sensitivity throughout the light exposure using a melanopsin-only model.

| Time | Young |  |  | Older |  |  |
| --- | --- | --- | --- | --- | --- | --- |
| | $\lambda_{\max}$ | $R^2$ | $Adj. R^2$ | | $R^2$ | $Adj. R^2$ |
| 15' | 485.3 | 0.66 | 0.62 | 495.3 | -0.39 | -0.59 |
| 30' | 485.3 | 0.79 | 0.76 | 495.3 | 0.52 | 0.45 |
| 45' | 485.3 | 0.95 | 0.94 | 495.3 | 0.48 | 0.41 |
| 60' | 485.3 | 0.93 | 0.92 | 495.3 | 0.64 | 0.59 |

Predicted peak spectral sensitivity using the melanopsin-only model during at 15-min intervals in young and older participants.

**Approach 2:** We then evaluated the combined contribution of melanopsin and cone photoreceptors in the melatonin suppression using a backward stepwise selection approach (Zuur et al., 2009). First, the Govardovskii alpha-band template (Govardovskii et al., 2000) was used to fit melanopsin as well as S-, M-, and L-cone opsin nomograms on the average, control-adjusted, percentage melatonin suppression, measured at each of the four time-intervals of light exposure in the young and older groups. The amplitudes assigned to each photoreceptor, *a1*, *a2*, *a3*, *a4,* were not constrained but allowed to be optimally determined by the nonlinear least-squares curve-fitting procedure (Eq. 4).

Peak sensitivities were kept at 420 nm for S-opsin (S-cones), 530 nm for M-opsin (M-cones), and 558 nm for L-opsin (L-cones) (CIE, 2018). Melanopsin sensitivity was fixed at 480 nm (CIE, 2018).

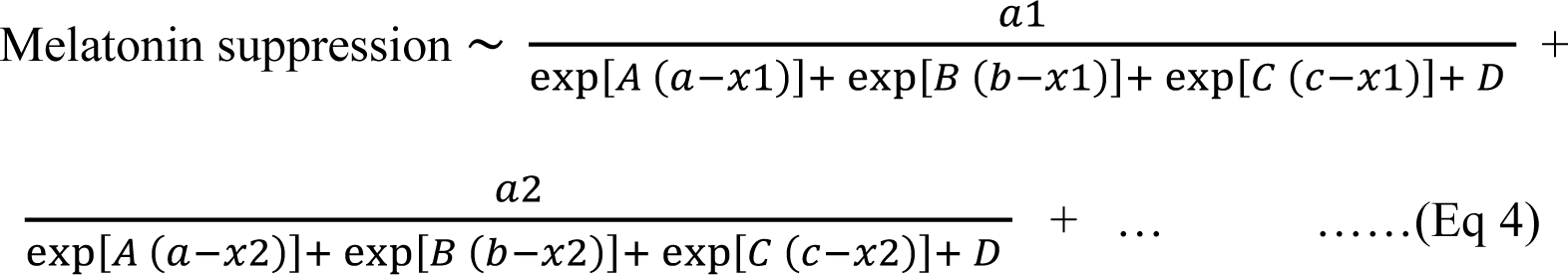

where 𝑥1 = 480/*wavelength,* 𝑥2 = 420/*wavelength* and so on. Other parameter values are the same as in Eq 3.

Thus for a particular time interval, melanopsin nomogram with amplitude *a1*, and cone opsin nomograms (S-, M-, L-cones, with amplitude *a2*, *a3*, *a4,* respectively) were simultaneously fitted on the melatonin suppression data. S-cones contribution was constrained to be of negative amplitude in accordance with Spitschan et al., (2014), Dacey et al., (2005), Woelders et al., (2018), while melanopsin, M- and L-cones contribution were not constrained; their amplitude could be either positive or negative, as determined by the modelling procedure. As in the first approach, we corrected for ocular media transmittance. The two lens filtering spectra obtained in the younger and older age groups (Najjar et al., 2014) were applied as a multiplicative term to each sensitivity functions.

We then explored photoreceptoral contributions in our melatonin suppression data from this initial candidate model of four photoreceptors (melanopsin - S + M + L-cones). The amplitude parameter for each photoreceptor nomogram (*a1*, *a2*, *a3*, *a4*) was estimated by the nonlinear least-squares regression model in R. The *P*-value of the t-test carried out to predict the significance of each amplitude parameter was determined, with significant level set at p < 0.05. The least significant parameter was dropped and the model refitted at each successive step, in a backward stepwise selection approach (Zuur et al., 2009). Thus, we began an iterative process whereby the photoreceptor showing the lowest unsignificant contribution was iteratively dropped from the initial four photoreceptor model, until significant photopigment contribution(s) was detected, if any (see Supplementary Tables 1 and 2). We thus evaluated the photopigment(s) contributing to melatonin suppression by light for each group and at each time interval, allowing to propose a model for the dynamics of melatonin suppression in the young and the older, at each respective duration of light exposure. If more than one photoreceptor was involved (e.g., melanopsin + L-cones), the fitted functions were thus a mathematical combination of photoreceptor nomograms accounted for pre-receptoral filtering by multiplying the function with its respective ocular lens transmittance spectra (Najjar et al., 2014). We assessed the assumptions of normality (Shapiro-Wilk’s test), homogeneity (Levene’s test), and independence (lag plot) of the residuals in order to validate each optimal model (Field et al., 2012). These assumptions were not violated, validating the predicted optimal models of photoreceptor contribution at each time interval and for each age group.

## Results

### Amplitude and phase of melatonin secretion in both groups

All participants displayed a significant nocturnal secretion of melatonin. The amplitudes of melatonin secretion assessed during the control session, were not different between young (65.8 ± 20.9 pg/ml) and older participants (89.3 ± 27.3 pg/ml) (*P* = 0.5). DLMOs, as markers of circadian phases, were not different between groups (*F* (1, 11) = 1.36, *P* = 0.26) and did not vary across night sessions (*F* (9, 99) = 0.43, *P* = 0.92). Mean DLMO was 2201 ± 0027 h in the young and 2228 ± 0028 h in the older participants.

### Melanopsin alone drives melatonin suppression by light in young but not older participants

Given that melanopsin is the predominant driver of melatonin suppression (Prayag et al., 2019), we evaluated melanopsin-only contribution in the response levels we observed. Our approach was further supported by recent results demonstrating that melanopsin fully accounts for the different non-visual responses observed across a wide range of light stimuli (Brown, 2020). In young participants, the fitted melanopsin-only nomograms at the four time intervals yielded adjusted R² values ranging between 0.62 - 0.94 (Figure 1A-D, Table 1). In older participants, adjusted R² values were systematically lower compared to those in the young, ranging from -0.59 to 0.59 (Figure 1E-H; Table 1). The negative adjusted R² value (−0.59) at 15 min in the older is indicative of a poor regression and is therefore not an appropriate model of melatonin suppression. Peak melatonin suppression was at ∼ 485 nm at all time intervals in the young group (Figure 2A; Table 1). In the older group, peak melatonin suppression was at ∼495 nm at all time intervals (Figure 2B; Table 1). These disparities between the two groups, i.e., the shift in towards the green part of the visible spectrum by ∼10 nm in the older group, as well as the poorer R² values, are consistent with a multi-opsin contribution to melatonin suppression by light in the older participants.

**Figure 1.**
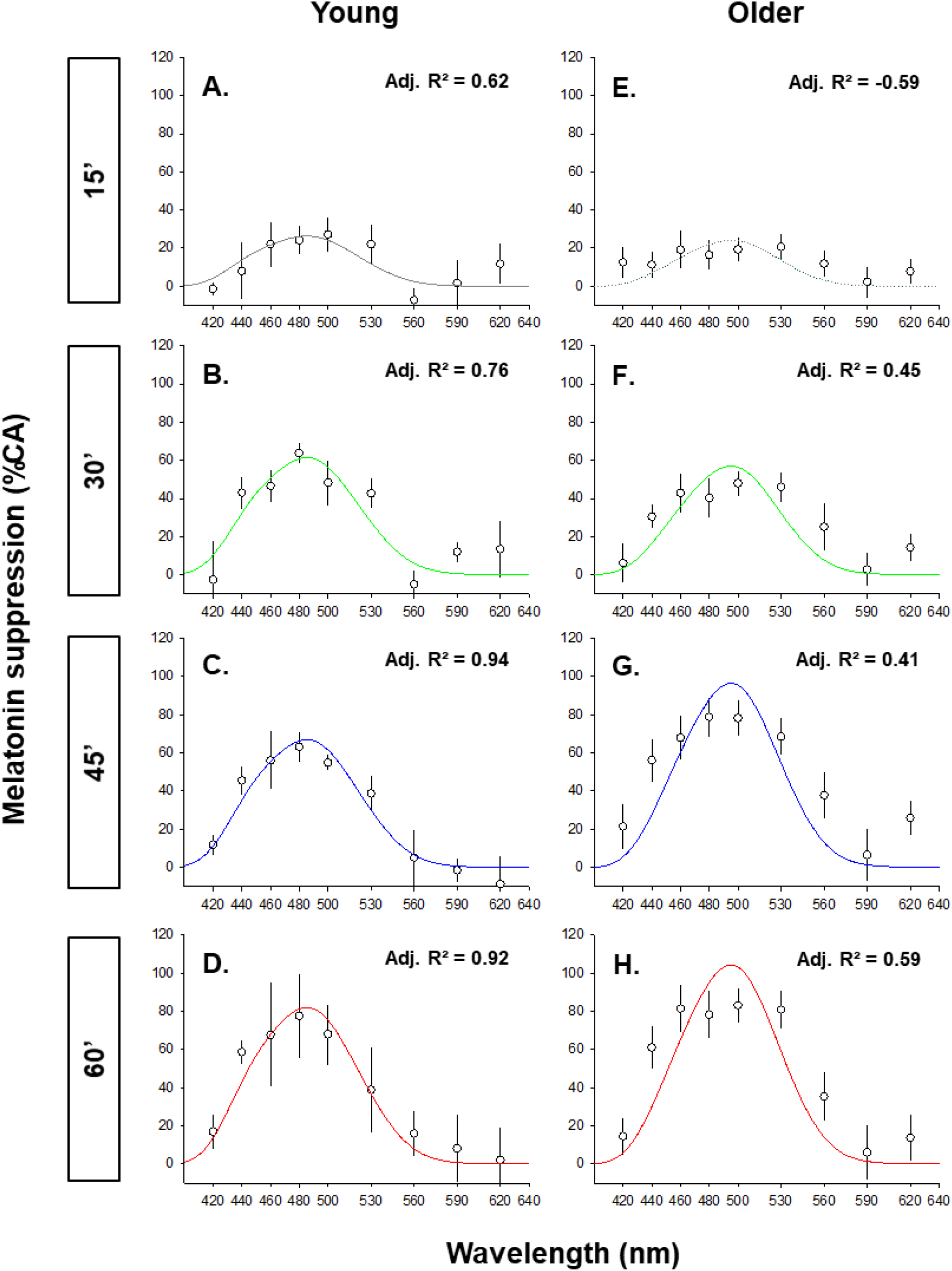
The dynamic spectral sensitivity of melatonin suppression by light in young and older participants, fitted with a melanopsin-only model. A melanopsin sensitivity function was fitted on the average, control-adjusted, percentage melatonin suppression, measured at each time interval for the young (A-D) and older (E-H) participants. Quantified melatonin suppression values are shown in open circle as average ± S.E. Each melanopsin-only nomogram was corrected for the respective transmittance of the crystalline lens. The adjusted R² of the model at 15 min in older participants was negative, indicating a poor/unreliable fit represented by a dotted grey line. The adjusted R² for each fitted functions is given as a measure of the goodness of fit and for comparison with literature.

**Figure 2.**
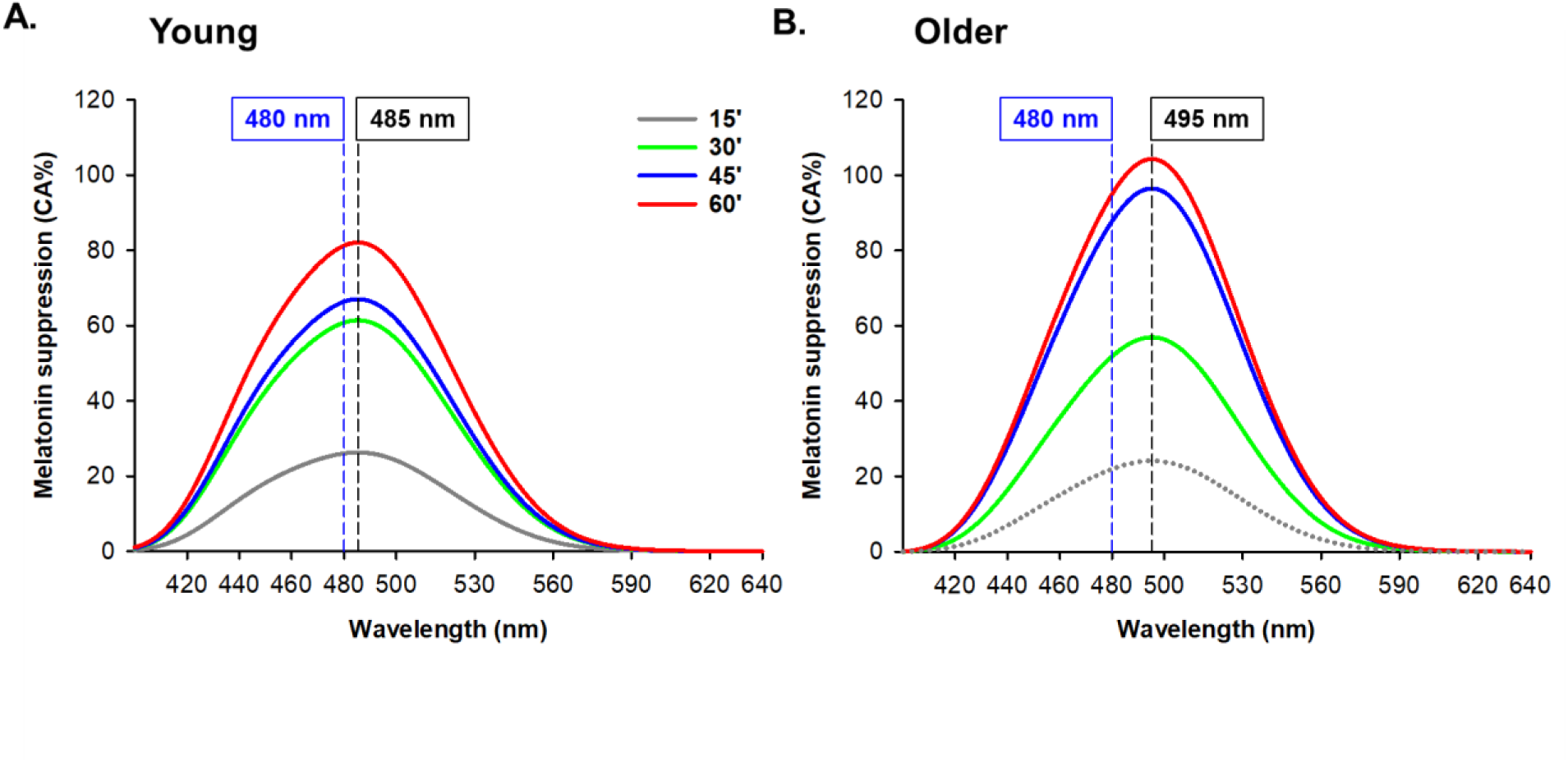
Overlaid spectral sensitivity curves (melanopsin-only model) for melatonin suppression by light in young and older participants throughout the 60 min light exposure. Overlaid spectral sensitivity curves (corrected for the respective transmittance of the lens) obtained in Figure 1. **A.** The melanopsin-only fitted function predicts a constant peak spectral sensitivity at ∼485 nm (dashed vertical line) in young participants over time, at 15, 30, 45 and 60-min of light exposure. **B**. The melanopsin-only fitted function predicts a constant peak spectral sensitivity different from the young by 10 nm, at ∼495 nm (dashed vertical line) in older participants after 15, 30, 45 and 60-min of light exposure. The adjusted R² at 15 min in older participants was negative, indicating a poor/unreliable fit represented by a dotted grey line. The dashed vertical line at 480 nm represents the peak spectral sensitivity of melanopsin. The predicted peak in each group and R² at each 15-min interval are provided in Table 1.

### Melanopsin + L-cones drive melatonin suppression by light in the older participants

We evaluated the combination of the four retinal photoreceptors (melanopsin, S-, M-, L-cones) driving the suppression response. Dropping the least significant contribution for each group and each interval (Supplementary Table 1-2), the best predictor of melatonin suppression in the young group, at all of the time intervals, was melanopsin with and R² values are identical to the melanopsin-only function (Figure 3A-D; Table 2). Peak melatonin suppression remained stable at ∼485 nm in the young group across all time intervals (Figure 4A). Conversely, in the older group, the model that fitted our data best was the melanopsin + L-cone model, at the 30, 45, 60-min intervals (Figure 3E-H; Supplementary Table 2) with respective adjusted R² values of 0.61, 0.51 and 0.63 (Table 2). At these time intervals, R² values of the melanopsin + L-cone model were higher than those of the melanopsin-only model (Table 1, Table 2), indicating better modelling of the suppression by the melanopsin + L-cone model in the older group. Again, the negative adjusted R² at 15 min in the older group (−0.26) is indicative of a poor regression that is therefore not an appropriate model of melatonin suppression. In the older group, for the melanopsin + L-cone model, an appropriate fit was first found at 30 min of light exposure and remained relatively constant and with a peak sensitivity in the range of 497.6 to 498.9 nm for the entire duration of the light exposure. (Figure 4B).

**Figure 3.**
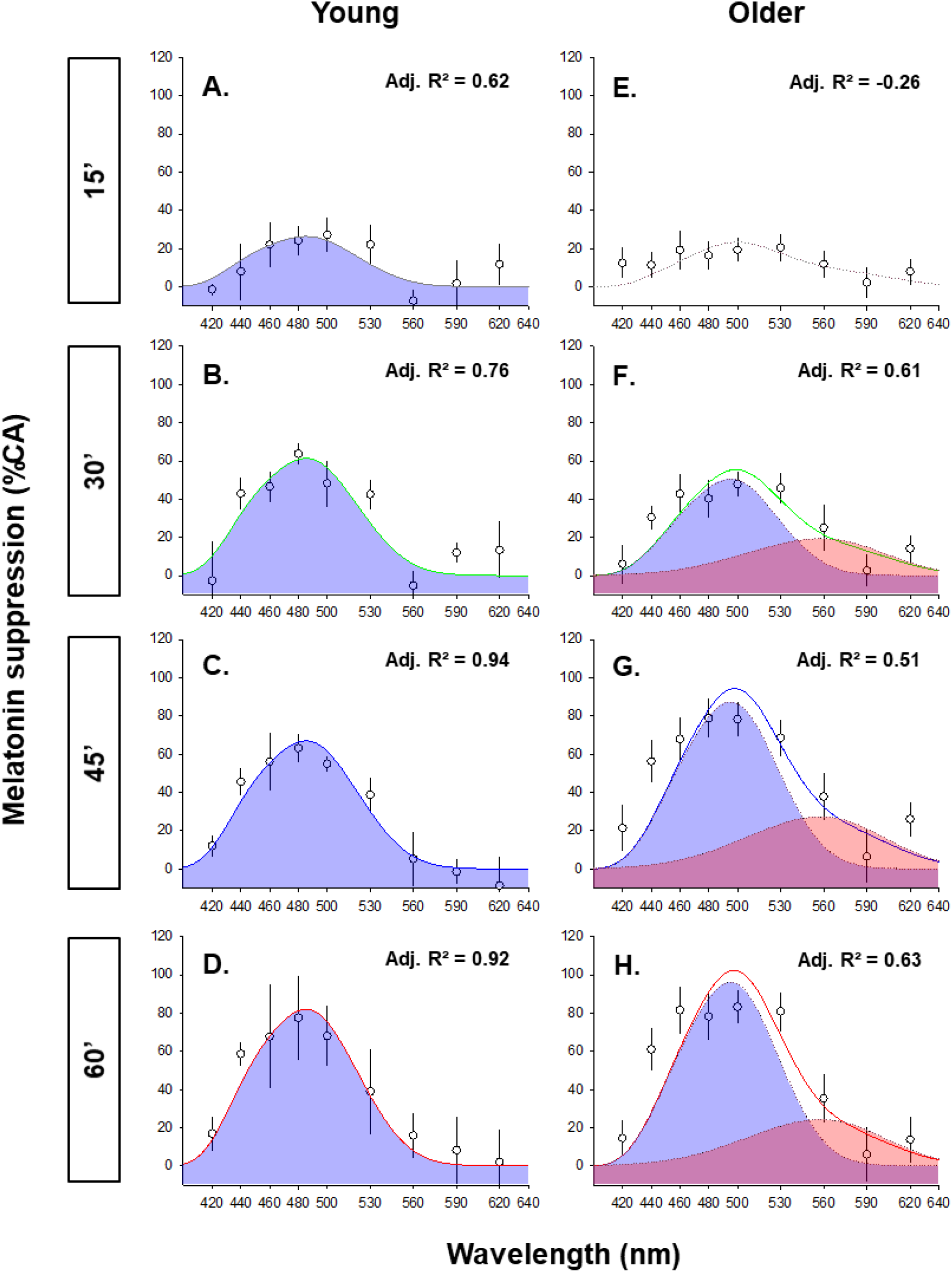
The dynamic spectral sensitivity of melatonin suppression by light in young and older participants fitted with a composite (melanopsin + L-cones) model. **A.** Spectral melatonin suppression by light in the young was robustly predicted using a melanopsin-only model at the 15, 30, 45, and 60 minute time points of the light exposure. As expected, the peak spectral sensitivity and R² values were identical to the melanopsin-only function fitted in Figure 1A. **B.** In older participants, our modelling predicted significant melanopsin + L-cones contribution to the spectral melatonin suppression response. Thus, the fitted function (corrected for the transmittance of the lens) was a mathematical combination of melanopsin (blue shaded area) and L-cones (red shaded area) nomograms at 30, 45, 60 min. Quantified melatonin suppression values are shown in open circle as average ± S.E. Each melanopsin-only nomogram was corrected for the respective transmittance of the crystalline lens. The adjusted R² for each fitted functions is given as a measure of the goodness of fit and for comparison with literature.

**Table 2:** Peak spectral sensitivity throughout the light exposure using a melanopsin + L-cones model.

| Time | Young |  |  | Older |  |  |
| --- | --- | --- | --- | --- | --- | --- |
| | $\lambda_{\max}$ | $R^2$ | $Adj. R^2$ | | $R^2$ | $Adj. R^2$ |
| 15' | 485.3 | 0.66 | 0.62 | 500.1 | 0.05 | -0.26 |
| 30' | 485.3 | 0.79 | 0.76 | 498.9 | 0.70 | 0.61 |
| 45' | 485.3 | 0.95 | 0.94 | 498.1 | 0.63 | 0.51 |
| 60' | 485.3 | 0.93 | 0.92 | 497.6 | 0.72 | 0.63 |

**Figure 4.**
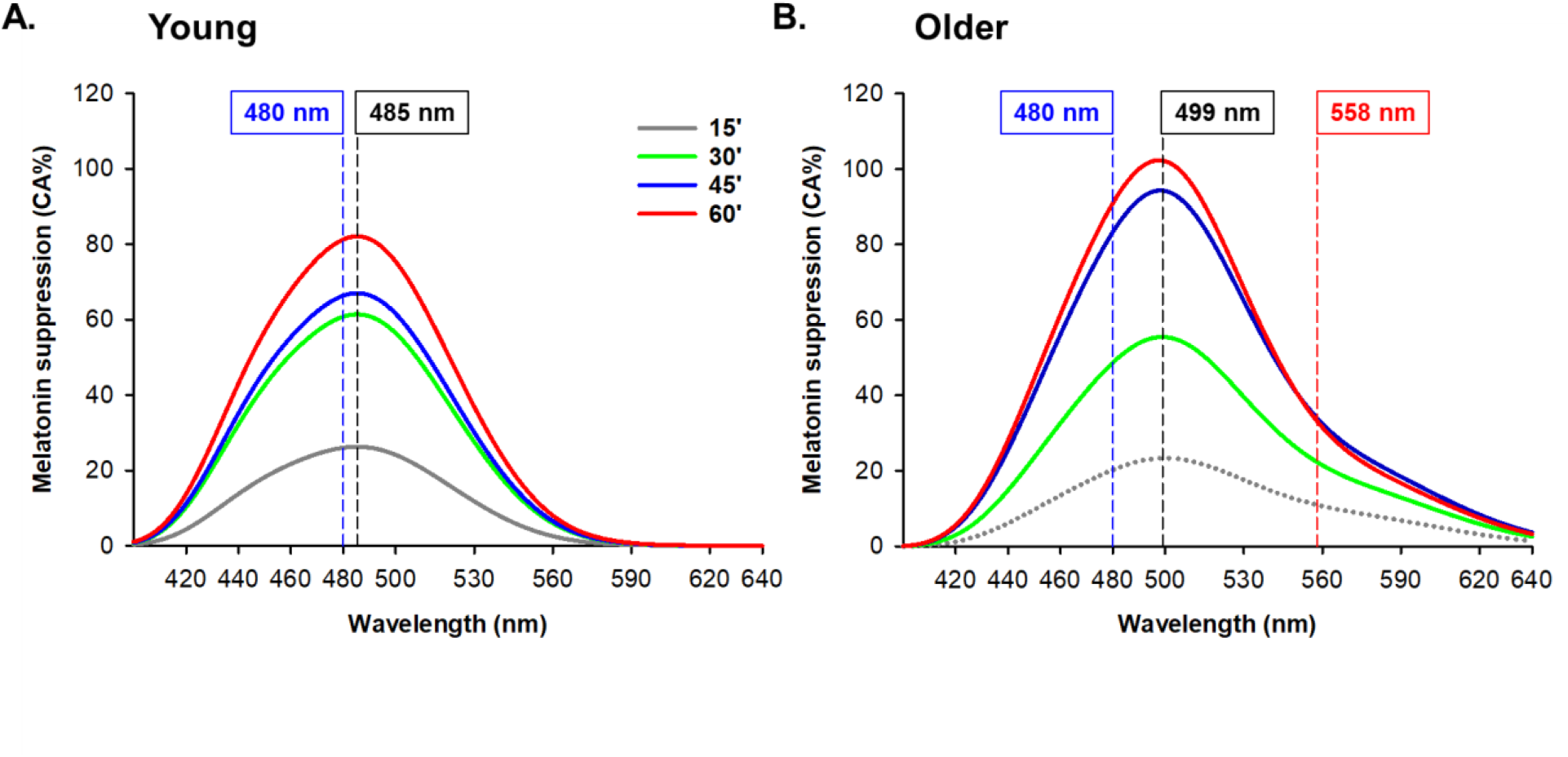
Overlaid spectral sensitivity curves (composite melanopsin + L-cones model) for melatonin suppression by light in young and older participants throughout the 60-minute light exposure. Overlaid spectral sensitivity curves (corrected for the respective transmittance of the lens) obtained in Figure 3. **A.** In young participants, spectral melatonin suppression was robustly predicted using a melanopsin-only model at 15, 30, 45, and 60-min intervals respectively. Dashed vertical lines at 485 nm represent the constant peak spectral sensitivity over time in the young group. **B.** In older participants, spectral melatonin suppression was better predicted by a composite melanopsin + L-cone model at the 30, 45, and 60 minute time points of the light exposure. Dashed vertical line at 499 nm represent the predicted average peak spectral sensitivityover time of the combined melanopsin + L-cones function in the older group. The predicted peak in each group and R² at each 15-min interval are given in Table 2. The dashed vertical lines at 480 nm and 558 nm represent the peak spectral sensitivity of melanopsin and L-cones, respectively.

## Discussion

Our results show that although melatonin suppression by light is predominantly, if not solely, driven by melanopsin in young individuals, a melanopsin-based model is insufficient to characterize melatonin suppression by light in older individuals. While melanopsin input remains predominant in the elderly, it is coupled with the contribution of L-cones leading to a shift in peak non-visual spectral sensitivity from ∼485 nm to ∼499 nm. Interestingly, these peak sensitivities and photoreceptor contributions did not change over the course of a 60-minute light exposure in the young and older. In young individuals, our model estimated a peak for melatonin suppression at ∼485 nm after 15 minutes of monochromatic light. This peak remained constant throughout the 60-minute light exposure. This spectral sensitivity of melatonin suppression, is close to the 480 nm peak sensitivity of melanopsin (Berson et al., 2002; Dacey et al., 2005) and therefore supports the predominant role of melanopsin in melatonin suppression during the 60-minute light exposure. On the other hand, in older individuals, 15 minutes of exposure to narrow-band light was not sufficient to induce a reliable spectral sensitivity curve (*i.e.,* with a positive adjusted R² value). Hypothetically, this may be due to a reduced non-visual sensitivity to short duration light exposures (i.e., 15 min) in the elderly. The peak spectral sensitivity of melatonin suppression, reliably established only after 30 minutes of light exposure, was relatively stable around 498-499 nm across the 60 minutes of light exposure. These results suggest that an additional opsin, besides melanopsin, is involved in melatonin suppression in older individuals. At all time-intervals, only an additive contribution of melanopsin and L-cones (+melanopsin+L) can fully describe the spectral melatonin suppression in the elderly. In addition, L-cones’ drive seems to increase over the duration of the light exposure. In older participants, our model suggests that a 558 nm light yields 22, 34, and 33% melatonin suppression at 30, 45 and 60 min, respectively compared to 3, 7, 7 and 9% in the young at 15, 30, 45 and 60 min, respectively (Figure 4). Altogether, this coupled melanopsin and L-cone additive input, strongly explains the shift in peak spectral sensitivity of melatonin suppression from 485 nm in the young to ∼499 nm in the older, and is in accordance with data showing excitatory L-cone input to ipRGCs in primates (Dacey et al., 2005). A similar additive input of melanopsin and L-cone has also been observed for the PLR (Spitschan et al., 2014; Woelders et al., 2018; Spitschan, 2019).

We suggest that retinal photoreceptor adaptation, consistent with the reorganisation that the human retinal network undergoes with aging (Eliasieh et al., 2007), is one possible mechanism behind the differences in photoreceptor contribution between young and older group. They might also result from adaptation mechanisms of the non-visual response *per se*. For instance, Giménez et al., (2014) showed that melatonin suppression was lower in subjects wearing blue-blocking contact lenses during a 30-min light exposure. However, after 16 days of continuously wearing blue-blocking lens, melatonin suppression was not different anymore from the control condition (normal contact lenses) in young healthy adults. These findings, suggest that non-visual photoreception is plastic and can undergo quick adaptations/compensations in young individuals. Similar compensatory mechanisms can also occur in older individuals who have less blue light reaching the retina due to ocular lens yellowing (Norren and Vos, 1974; van de Kraats and van Norren, 2007; Najjar et al., 2016). In fact, Najjar et al., (2014)have shown that although lens yellowing leads to a 42% decrease in short-wavelength light reaching the retina in the older individuals, this was not associated with an attenuation of melatonin suppression to short-wavelength lights. Taken together, retinal restructuring and reorganisation coupled with potential adaptation/compensatory mechanisms (Eliasieh et al., 2007; Gollisch and Meister, 2010; Samuel et al., 2011), could explain age-related changes in photoreceptoral contribution to non-visual responses shown in our study.

Findings from this study confirm the peak spectral sensitivities observed by Najjar et al., (2014) following 60 minutes of light in young and old participants and corroborate melanopsin stimulation by light as the predominant predictor of non-visual responses. Of note, Brown (2020) recently observed that the degree melanopsin stimulation, calculated via melanopic illuminances, accurately predicts the response levels of melatonin suppression and circadian phase resetting, across an array of lighting types, light spectra and exposure durations. In that study, analysis was restricted to an age range of 20 to 45 years, which would therefore include our young group (mean age of 25.8) but not our old group (mean age of 59.4). Consistent with Brown, (2020), our results in young participants confirm the observation that melanopsin can solely account for melatonin suppression. They are also consistent with results by Prayag et al., (2019), which show that melanopsin is the predominant contributor to melatonin suppression in response to a 90 minutes of monochromatic light exposure in an equivalent young age group (mean age of 24.5 years). Our results in the older individuals do not invalidate the conclusions of these (Prayag et al., 2019; Brown, 2020) but indicate that the relative contribution of retinal photoreceptors is dependent on the non-visual response measured, the duration and irradiance of the light exposure (Lucas *et al*. 2014; McDougal & Gamlin 2010; Spitschan *et al*. 2014), and is likely to vary with age.

Our results do not show significant S-cones contribution to the acute melatonin suppression response, both in young and older participants. This is in agreement with recent findings showing no evidence of S-cones involvement in melatonin suppression following differential S-cone excitation using silent substitution (Spitschan et al., 2019a). Nevertheless, previous studies have reported a peak sensitivity for melatonin suppression at ∼460 nm (Brainard et al., 2001, 2008; Thapan et al., 2001), which is at a proximity to the midpoint between the peak sensitivities of S-cones (∼420 nm) and melanopsin (∼480 nm) (Dartnall et al., 1983; Spitschan et al., 2019b). Although these findings suggest an additive input from both photopigments to the acute light-driven melatonin suppression response, our data and models do not support any S-cones contribution to melatonin suppression in young and older individuals, across a 60-minute light exposure.

### Limitations

Our study has a few limitations. First, the sample sizes in both groups were small (n= 5 in the young group, n= 8 in the older). While independent, single measure comparisons between both groups would have been insufficient, our protocol is a repeated-measure within-subject design which assessed each participant repeatedly, and randomly, over 10 experimental conditions, which equates to 130 individual data points. We also implemented a fitting procedure to estimate the peak sensitivity as well as a backward stepwise selection procedure to investigate photoreceptor contribution. The determination coefficients found in the two steps of our analysis, both in the young and old, are indicative of high effect sizes and statistical power, and provide confidence in the differences we observed. Second, given the proximity of the peak sensitivity between M- and L-cones (530 nm vs. 558 nm; Dartnall et al., 1983; Spitschan et al., 2019b), we cannot exclude a potential M-cones input in the suppression response in the older. This does not invalidate our results of an additive combination of photoreceptors, melanopsin and L-cones, in the older. Finally, we also recognize that the photoreceptoral contribution may yet differ across non-visual responses to light. Thus, the photosensitivity predictions of our models only apply for the acute melatonin suppression response which, we further stress, is not a reliable proxy for SCN-dependent circadian responses in humans (Rahman et al., 2018).

## Conclusion

Our study highlights unexpected yet important physiological traits of the aging of non-visual photoreception and its impact on melatonin suppression. While melanopsin contribution is predominant in both young and older, a robust input from L-cones contributes to melatonin suppression by light in older participants. These results carry an important translational value as they suggest that artificial lighting used in clinical settings, for instance for light therapy of sleep or affective disorders, or even in everyday life (e.g., blue blocking filters) may not show the same efficiency in older patients.

## Acknowledgements

This analytical work was supported by grants 12-TECS-0013-01 and 16-IDEX-0005 from ANR (Agence Nationale de la Recherche) to C.G., a grant from GIS Aging to C.G., by a Doctoral Fellowship from the French Ministry of High Education and Research to R.P.N and the ASPIRE-NUS startup grant (NUHSRO/2022/038/Startup/08) to R.P.N.. The funders had no role in study design, data collection and analysis, decision to publish, or preparation of the manuscript. The authors wish to thank H.M. Cooper and C. Chiquet for respectively funding (EU-FP7 (EUCLOCK)) and conducting some of the experimental sessions in the young for in the initial study (Najjar et al 2014); B. Claustrat for the melatonin assays; P-L Cornut for the ophthalmologic examination of the participants; and P. Denis for the use of the experimental facility (Hopital Edouard Herriot, Lyon, France).

## Data availability

The raw data supporting the conclusions of this manuscript will be made available by the authors, without undue reservation, to any qualified researcher.

## Author contributions

The analyses described in this article were conceived, designed and performed by R.P.N., A.S.P., and C.G.; R.P.N., A.S.P., and C.G interpreted the data and wrote the manuscript.

