## Supplementary Information for "Melatonin suppression by light involves different retinal photoreceptors in young and older adults"

### Supplementary methods

#### Modelling approach evaluating photoreceptor contribution

The first row in supplementary tables 1 and 2 is our starting model, a combination of four retinal photoreceptors: melanopsin - S+ M + L-cones. We explored photoreceptoral contribution in our melatonin suppression data from this initial candidate model of four photoreceptors (melanopsin - S + M + L-cones). The next rows are associated with a photoreceptor term being dropped. The *P*-value of the t-test carried out to predict the significance of each amplitude parameter was determined, with significant level set at $p < 0.05$. The least significant parameter was dropped and the model refitted at each successive step without this photoreceptor contribution, in a backward stepwise selection approach (Zuur et al., 2009). Thus, we began an iterative process whereby the photoreceptor showing the lowest unsignificant contribution was iteratively dropped from the initial four photoreceptors model, until significant photopigment contribution(s) was/were detected. The last row is our final model, which is the photoreceptor (s) best predicting the melatonin suppression in each group and each time interval.

### Supplementary tables

**Supplementary Table 1.** Photoreceptor contribution to melatonin suppression in young participants

| | | ***Nomogram fit significance*** **within the model** *(p-value)* | | | |
|---|---|---|---|---|---|
| | **Models** | **Melanopsin** | **S-cones** | **M-cones** | **L-cones** |
| ***15'*** | melanopsin - s + m + l | 0.023 | 0.33 | 0.67 | 0.69 |
| | melanopsin - s + m | 0.005 | 0.37 | 0.88 | - |
| | melanopsin - s | 8.53E-05 | 0.36 | - | - |
| | **melanopsin only** | **1.78E-05** | - | - | - |
| ***30'*** | melanopsin - s + m + l | 3.19E-04 | 0.85 | 0.44 | 0.35 |
| | melanopsin + m + l | 3.81E-06 | - | 0.44 | 0.35 |
| | melanopsin + l | 2.86E-10 | - | - | 0.56 |
| | **melanopsin only** | **3.19E-12** | - | - | - |
| ***45'*** | melanopsin - s + m + l | 2.28E-04 | 1.00 | 0.64 | 0.59 |
| | melanopsin + m + l | 1.23E-06 | - | 0.60 | 0.56 |
| | melanopsin + l | 1.26E-13 | - | - | 0.78 |
| | **melanopsin only** | **2.38E-15** | - | - | - |
| ***60'*** | melanopsin - s + m + l | 0.002 | 1.00 | 0.65 | 0.54 |
| | melanopsin + m + l | 2.72E-05 | - | 0.61 | 0.51 |
| | melanopsin + l | 2.03E-09 | - | - | 0.60 |
| | **melanopsin only** | **2.93E-11** | - | - | - |

**Supplementary Table 2.** Photoreceptor contribution to melatonin suppression in older participants

| | | ***Nomogram fit significance* within the model** *(p-value)* | | | |
|---|---|---|---|---|---|
| | **Models** | **Melanopsin** | **S-cones** | **M-cones** | **L-cones** |
| ***15'*** | melanopsin - s + m + l | *No reliable models (negative $R^2$)* | | | |
| ***30'*** | melanopsin - s + m + l | 7.78E-04 | 1.00 | 0.85 | 0.56 |
| | melanopsin + m + l | 1.46E-05 | - | 0.84 | 0.53 |
| | **melanopsin + l** | **2.39E-11** | - | - | **0.025** |
| ***45'*** | melanopsin - s + m + l | 1.61E-06 | 1.00 | 0.58 | 0.17 |
| | melanopsin + m + l | 1.52E-09 | - | 0.54 | 0.15 |
| | **melanopsin + l** | **2.87E-16** | **-** | **-** | **0.015** |
| ***60'*** | melanopsin - s + m + l | 4.02E-06 | 1.00 | 0.91 | 0.55 |
| | melanopsin + m + l | 5.68E-09 | - | 0.90 | 0.53 |
| | **melanopsin + l** | **<2E-16** | - | - | **0.038** |
